# Die hard in Lake Bourget! The case of *Planktothrix rubescens*

**DOI:** 10.1101/2021.02.02.429300

**Authors:** Marthe Moiron, Frédéric Rimet, Cyrille Girel, Stéphan Jacquet

## Abstract

Blooms of *Planktothrix rubescens* have been recorded for 15 years in Lake Bourget (France), from 1995 to 2009. Then, the presence of this filamentous and toxic cyanobacterium became anecdotic between 2010 and 2015 and it was clearly thought that such a proliferation was over. However, against all odds, blooms occurred again in 2016 and 2017 despite apparent very low phosphorus concentrations in surface waters of the lake. Aims of this study were thus to explain the reasons of this come back in order to propose scenarios likely to be helpful to stakeholders who need to know if such proliferations may occur again in the future. We show that phosphorus input, both from the main tributaries to the lake and possibly from the sediments, were likely the triggers of the new development of the cyanobacterium since a minimum autumn/winter inoculum of *P. rubescens* was detected the year before. Then, the bloom, that was observed deeper than previous years, was associated to a conjunction of factors already well-known to favour the development of this very competitive species (i.e. mild winter temperature, water column stability, available light at depth, surface water transparency, low predation, etc…). Although many factors and processes could account for the occurrence and bloom of the cyanobacterium, not observed, measured or taken into account here, a plausible scenario could be proposed and may be useful to deciders. One thing remains unclear: where do the cyanobacterium hides when it is not observed during the routine monitoring survey and thus from which place it could initiate its development (nearshore, in the pelagic zone, from the sediment?), unless it is simply not sampled and observed due to methodological bias.

## Introduction

Lake Bourget, the largest deep natural lake in France, has undergone an eutrophication phase during the second part of the 20^th^ century, as many other ecosystems worldwide (Jenny et al. 2020). While the phosphorus concentration was lower than 10 µg/L before 1940, it reached up to 100 µg/L during the 1980s’ because of massive discharges from industrial effluents and insufficiently treated domestic wastewaters at that time (Jacquet et al. 2005). With eutrophication, biological changes were recorded, for instance within the eukaryotic microbial communities, firstly around 1940 and secondly between 1960 and 1980 (Capo et al. 2016, Capo et al. 2017). Management actions were set up in order to reduce nutrient concentrations in the lake, leading to a lower phosphorus concentration which dropped under 20 µg/L after 2000 (Capo et al. 2016, Capo et al. 2017, Jacquet et al. 2017). Significant ecological changes occurred in the pelagic compartments during the lake’s reoligotrophication, notably regarding the toxic filamentous cyanobacterium *Planktothrix rubescens* (Jacquet et al. 2014, Frossard et al. submitted).

Planktonic cyanobacterial species are characterized by a high level of adaptations and tolerances to various environments and can produce important amounts of toxins. Among these species, *P. rubescens* (Anagnostidis and Komarek 1988) is a red-coloured filamentous and gas-vacuolated cyanobacterium likely to develop in meso-to moderately eutrophic conditions (Jacquet et al. 2005, Dokulill and Teubner 2012). It is able to adjust via buoyancy its position in the water column to its preferred low light environment, and maybe in response to other resources, where it often forms deep water maxima in stratified lakes of temperate latitudes (Feuillade 1994, Micheletti et al. 1998, Bright and Walsby 2000, Vinçon-Leite et al. 2002, Zotina et al. 2003, Jacquet et al. 2005). This taxon is referred as an R-strategist according to Reynolds et al. (2002), and is known as an important hepatotoxic microcystins (MCs) producer (Fastner et al. 1999, Briand et al. 2005) likely to be harmful to a large variety of animals including human health (Sotton et al. 2011, 2012, Kurmayer et al. 2015). Indeed, *P. rubescens* produces a variety of microcystins, especially MC-LR and MC-RR, and these toxins have been shown to contaminate different fish tissues. Both filaments and toxins have been observed in intestinal tracts of whitefish and the presence of MC-LR has been detected in their intestine and liver. MCs were also detected in the muscles and liver of young perch of the year through dietary routes, particularly *via* the consumption of MC-containing *Daphnia* (Sotton et al 2011, 2012). Consequently, such toxins can be incorporated into different organisms by ingestion of *P. rubescens* filaments, leading to potential adverse effects on the animals’ health. The issue is particularly important since Lake Bourget is a place for an important professional and recreational fishing activity and is a source of drinking water for thousands of inhabitants.

While paleolimnological data first revealed the importance of this species in Lake Bourget during early years of eutrophication (i.e. >1930; Savichtcheva et al. 2015), *P. rubescens* was mainly observed and counted as a dominant species in the lake between the mid 1990’s and 2009. This was explained as a response to the reoligotrophication process of this lake, intermediate phosphorus levels, important water column stability, increasing transparency, etc… (Jacquet et al. 2005, 2014). The success of reoligotrophication was that from 2010, the species declined and “disappeared”, and, in the same time, phytoplankton biomass was considerably reduced, together with a change in taxon composition (Frossard et al. submitted). However, against all odds, regarding at first sight phosphorus concentration levels in surface waters of the lake, *P. rubescens* “reappeared” and important biomasses were recorded in 2016 and 2017.

This study aims at explaining this paradoxical situation (phosphorus depletion and *P. rubescens* reappearance) and proposing some scenarios to predict future possible developments of this cyanobacterium in Lake Bourget. Long-term ecological monitoring survey in this lake offers the possibility to analyze factors responsible for the development of cyanobacteria/algal species, therefore we used the dataset available for the lake and of its main tributaries in order to explain what could have been the reasons of the 2016-2017 bloom episodes of *P. rubescens*. Our hypotheses were that phosphorus input from rivers and/or sediment could be important factors triggering a new development of the cyanobacterium, whose growth and development could be sustained thereafter thanks to a conjunction of favorable conditions. Our results tend to confirm these hypotheses and highlight also the importance of a minimal autumn/winter concentration of the cyanobacterium (referred latter to as an inoculum) to develop further during the year.

## Methods

### Description of the site

Lake Bourget (45°44’N, 231 m altitude) is the largest natural deep lake in France and is located on the edge of the Alps. It is a warm, meromictic and elongated (18 and 3 km in length and width respectively), north-south orientated lake, with an area of 42×10^6^ m^2^, a total volume of 3.5×10^9^ m^3^, maximum and average depths of 145 m and 80 m respectively, and a water residence time of approximately 10 years. Winter overturn reaches the bottom of the lake only during very cold winters. It has a catchment area of about 560 km^2^, with maximum and average altitudes of 1,845 and 700 m respectively. There are two important cities beside the lake, Chambéry to the south and Aix-les-Bains to the east, with a combined population of 180,000 inhabitants, plus a large influx of tourists (>50,000) in summer. The lake suffered from eutrophication since 1950 and water quality restoration programs started in the 1970s. These programs involved the development and improvement of wastewater treatment plants, and in 1981, the diversion of the treated sewage of the two main cities from the lake, directly downstream the lake into the Rhône River. Other improvements of the functioning of the sewer system-especially the control of the combined sewer overflows-lead in the recent years to an additional reduction of the nutrient loading to the lake. Two rivers, the Leysse and the Sierroz, are the main inputs to the lake, with average flow rates of 6.5 and 2.5 m^3^/s, respectively. The flow rates of these two rivers can occasionally (during floods for instance) reach more than 120 and 30 m^3^/s, respectively. The outflow from the lake, located on its northern shore, is known as the Savière channel (length: 4.5 km, mean flow rate: 10-30 m^3^/s, annual output: 0.5 km^3^) and it flows into the Rhone River.

### Data

All the lake data used in this paper correspond to sampling performed at the reference sampling site located in the middle and deepest part of the lake, referred to as point B and are part of the lake observatory and its information system (Rimet et al. 2020, *© OLA-IS, AnaEE-France, IRLife, INRAE of Thonon-les-Bains, CISALB)*. This site is more than 1.5 km from each bank and more than 5 and 10 km from the Sierroz and Leysse rivers, respectively. A conductivity-temperature-depth (CTD) measuring device (CTD SEABIRD SBE 19 Seacat profiler) was used to obtain vertical profiles of temperature, dissolved oxygen, pH, and conductivity.

Temperature data were also used to determine the onset of water column stratification. We assumed that stratification had occurred when there was a temperature differential of more than 1°C between the 2- and 50-m depths on two consecutive sampling dates. The Brunt-Väisälä frequency (which measures the natural frequency of oscillation of a vertical column of water, and can be viewed as an index of the water column stability, Lemmin 1978) was also calculated from the temperature values, according to the following equation:

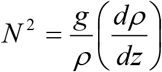

where: *N*^*2*^ is the stability coefficient (in s^-2^)

*g* is the acceleration parameter (in m/s^2^)

*ρ* is the water density (no unit)

*z* is the depth (in m)

with: *ρ*(T) = 1000 – 7×10^−3^ (T-4)^2^ according to Lemmin (1978)

where: T is the temperature (in °C)

Nutrient concentrations (such as total phosphorus and P-PO_4_, but also total nitrogen, N-NO_3_, N-NH_4_) were measured monthly to bi-monthly at ten different depths (from surface to bottom) according to normalised procedures and protocols (AFNOR 1982). The concentration of chlorophyll *a* was determined using the method of Strickland and Parsons (1972). Transparency data were obtained using a Secchi disk. Phytoplankton species and biovolumes were analysed according to the European standardised method (Afnor 2006) of Uthermöhl (1958). Species biovolumes used for this monitoring are available in Rimet & Druart (2018). For *P. rubescens*, the cell concentrations were estimated by counting 200-µm length filaments and by assuming a mean cell length of 5 µm. Several vertical profiles of the main phytoplankton groups, including *P. rubescens*, and of the temperature were also obtained using a submersible spectrofluorimeter (BBE-Fluoroprobe, Germany). This *in situ* measuring/recording device, which can be used to perform chlorophyll *a* analysis and integrated algal class determination, has been shown to provide a realistic estimation of the abundance and dynamics of the cyanobacterial population after specific calibration (Leboulanger et al. 2002).

River data come from two automatic sampling stations located in the two main tributaries, i.e. the Leysse and Sierroz, responsible for >75% of the water flowing into the lake. Samplers (ASP station 2000 Hendress + Hauser) are located at 1.3 km and 0.75 km from the lake, for the Leysse and the Sierroz, respectively. Waters for nutrient concentrations (such as total phosphorus, P-PO_4_, N-NO_3_ and N-NH_4_) are collected using a daily time scale and have been measured since 2003. This sampling allows obtaining an accurate estimation of the quantity of nutrient discharge into the lake as well as the key periods of such inputs.

Data such as air temperature, wind force and direction, irradiance, cloudiness, and precipitation were obtained at a 3-hour time step from the meteorological station Voglans at Chambéry airport, located less than 1 km from the southern shore of the lake.

## Results

### *Dynamics and distribution of* Planktothrix rubescens

*P. rubescens* began to bloom in Lake Bourget in 1995/1996. Each year until 2009 (except 2004), the cyanobacterium developed significantly reaching regularly >50% of the total phytoplankton biomass (Vinçon-Leite et al. 2002, Jacquet et al. 2005, 2014). In 2009, its cells concentration was still high, but much lower than in 2008 (a record year with 185,600 cells/mL recorded in July). However, following a conjunction of factors, as explained in Jacquet et al. (2014), the cyanobacterium “disappeared” during the winter 2009/2010 (Fig. 1A). However, at the end of 2015 (i.e. October-November) and during the 2015-2016 autumn/winter period, *P. rubescens* reappeared (Fig. 1B) and proliferated latter in the year 2016 reaching >50,000 cells/mL in September in the metalimnion, at a depth greater than observed by past (i.e. between 20 and 25 m *vs*. between 15 and 20 m for the period before 2009). In details, we observed at the end of October 2015 ∼500 cells/mL that were counted at 20 m and this concentration increased until mid-November to reach ∼2,000 cells/mL. During the following weeks of 2016, the cyanobacterium spread over the surface water column and remained observed along the year, reaching >15,000cells/mL in early summer and ∼50,000 cells/mL at the end of summer/early autumn. Then, a progressive decrease of the cyanobacterial biomass and a shift of *P. rubescens* cells towards the surface was recorded. Cells maintained however at a relatively high level (up to 8,000 cells/mL) at all depths between surface and 50 m during the winter 2016/2017 (Fig. 1B). Thereafter, in 2017, *P. rubescens* developed massively at depth (reaching >19,000 cells/mL by the end of May at 19 m, >22,000 cells/mL at 22.5 m on June 13, >31,000 cells/mL at 25 m on June 27 and >45,000 cells/mL at 21.5 m on July 10). High concentrations (>25,000 cells /mL) were measured until the end of September at various depths (Fig. 1B). During the winter 2017/2018 the cyanobacterium disappeared, and no new development has been observed until now, i.e. 2021 (Supplementary Fig. 1).

**Fig. 1.**
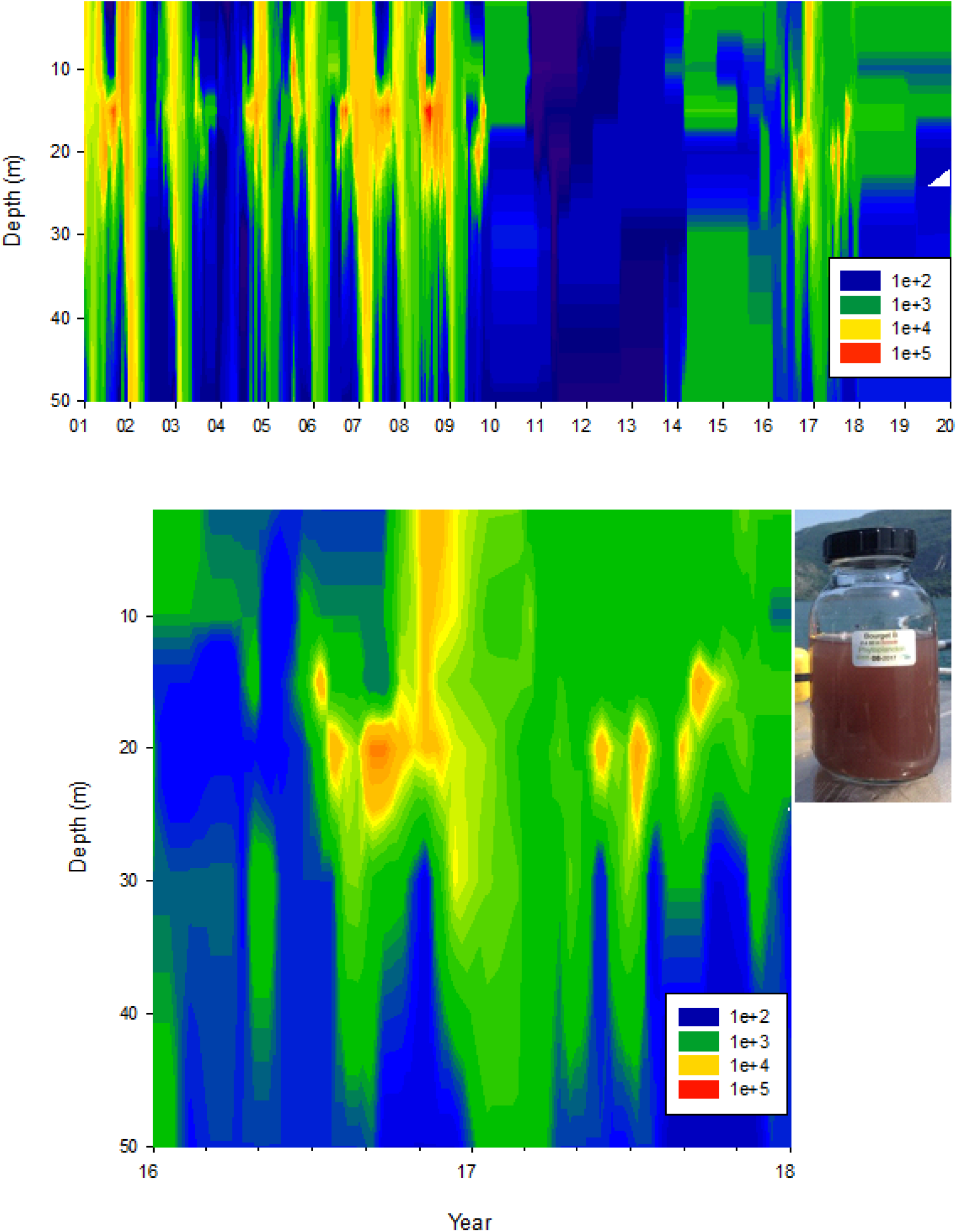
(A) *P. rubescens* dynamics and distribution from 2001 to 2019 at point B in Lake Bourget. (B) Zoom of the 2016-2017 bloom episode associated to a picture of a <62 µm plankton net sample obtained in spring 2017.

### Meteorological data

Over the period 2010-2019, the time of sunshine followed approximately the same annual pattern, with an increase during summer and a decrease in winter. While the summer sunshine level was almost similar each year, some differences were, however, recorded in winter. During winters 2012/13, 2017/18 and 2018/19, the daily time of sunshine stayed relatively low (< 4.0 h) from November to March. This phenomenon was even more important for the winter 2017-18 with a very low sunshine time (maximum and average sunshine time of 2.84 h and 2.17 h, respectively) and no significant fluctuation was recorded (Fig. 2A). By contrast, during winters 2015/16 and 2016/17 the sunshine decreased to low values but oscillated with higher values (Fig. 2B).

**Fig. 2a.**
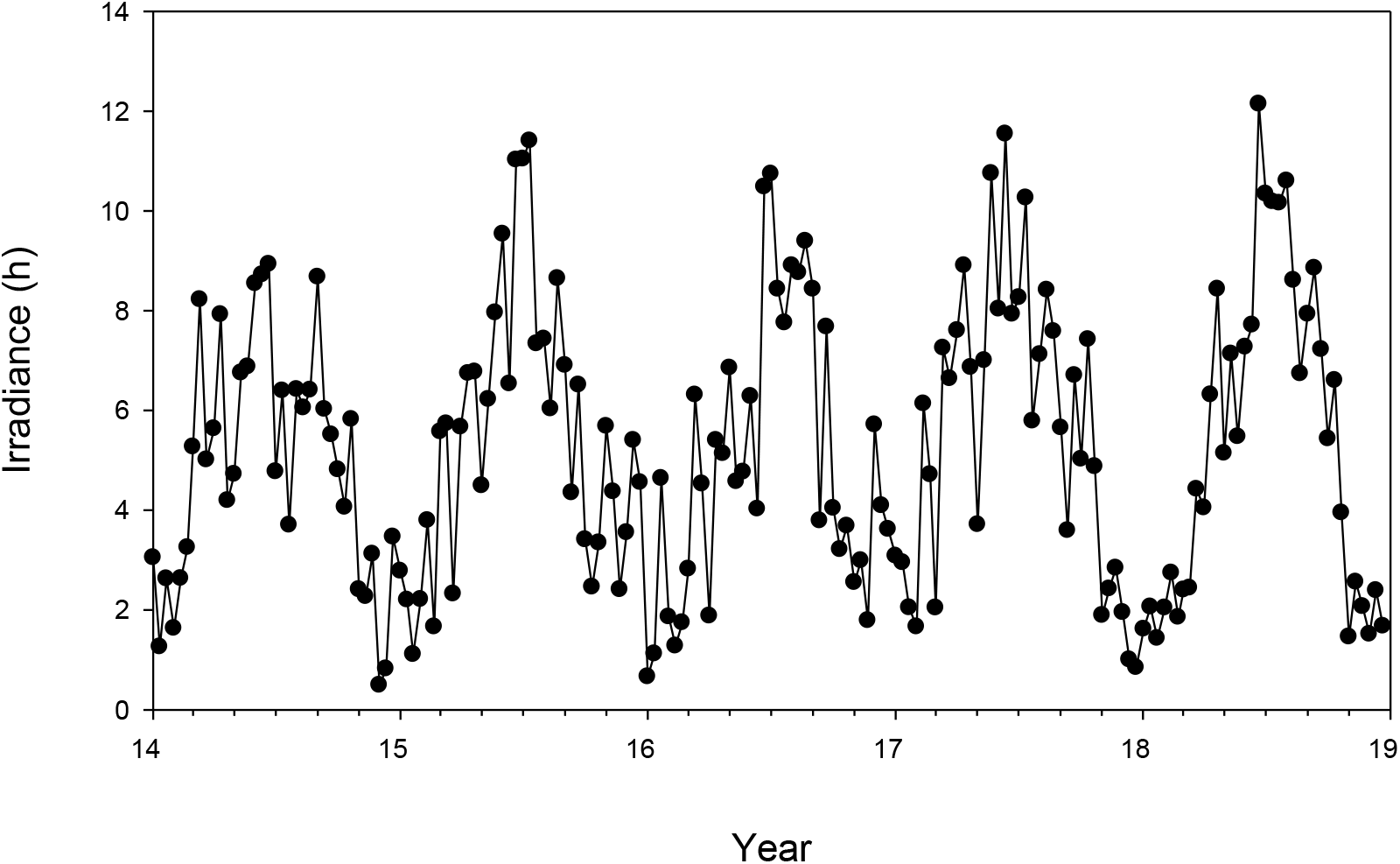
Chronicle of sunshine time averages from January 2014 to December 2018.

**Fig. 2b.**
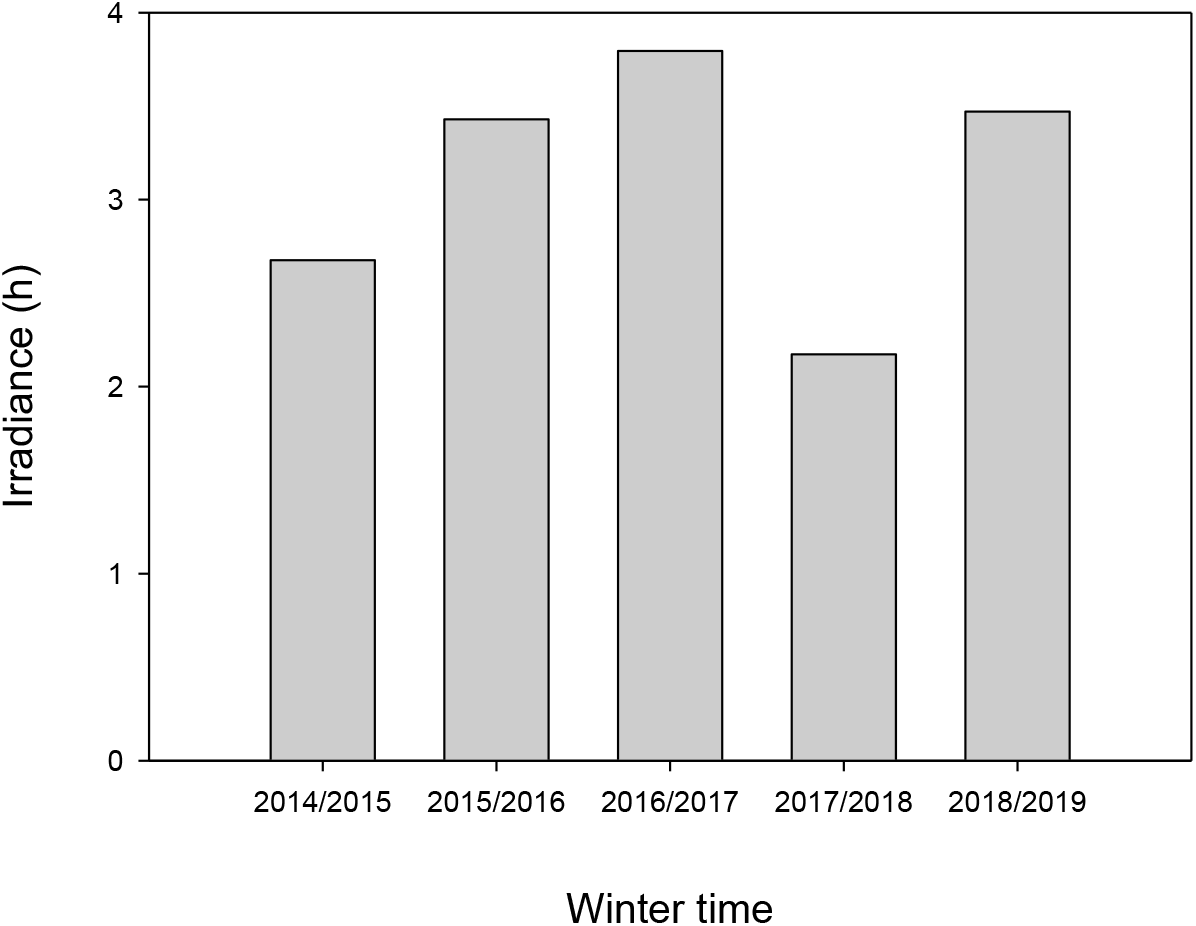
Average of sunshine time for winters 2014-15 to 2018-19.

### Transparency

The transparency of the water column ranged from 1.9 m to 14.4 m between 2010 and 2019. It was higher from 2010 to 2015 compared to subsequent years. Indeed, winter peaks were recorded with a maximum depth ranging from 13.0 m to 14.4 m, unlike the 2016-2019 period, where depths varied between 7.3 m and 10.6 m (Fig. 3).

**Fig. 3.**
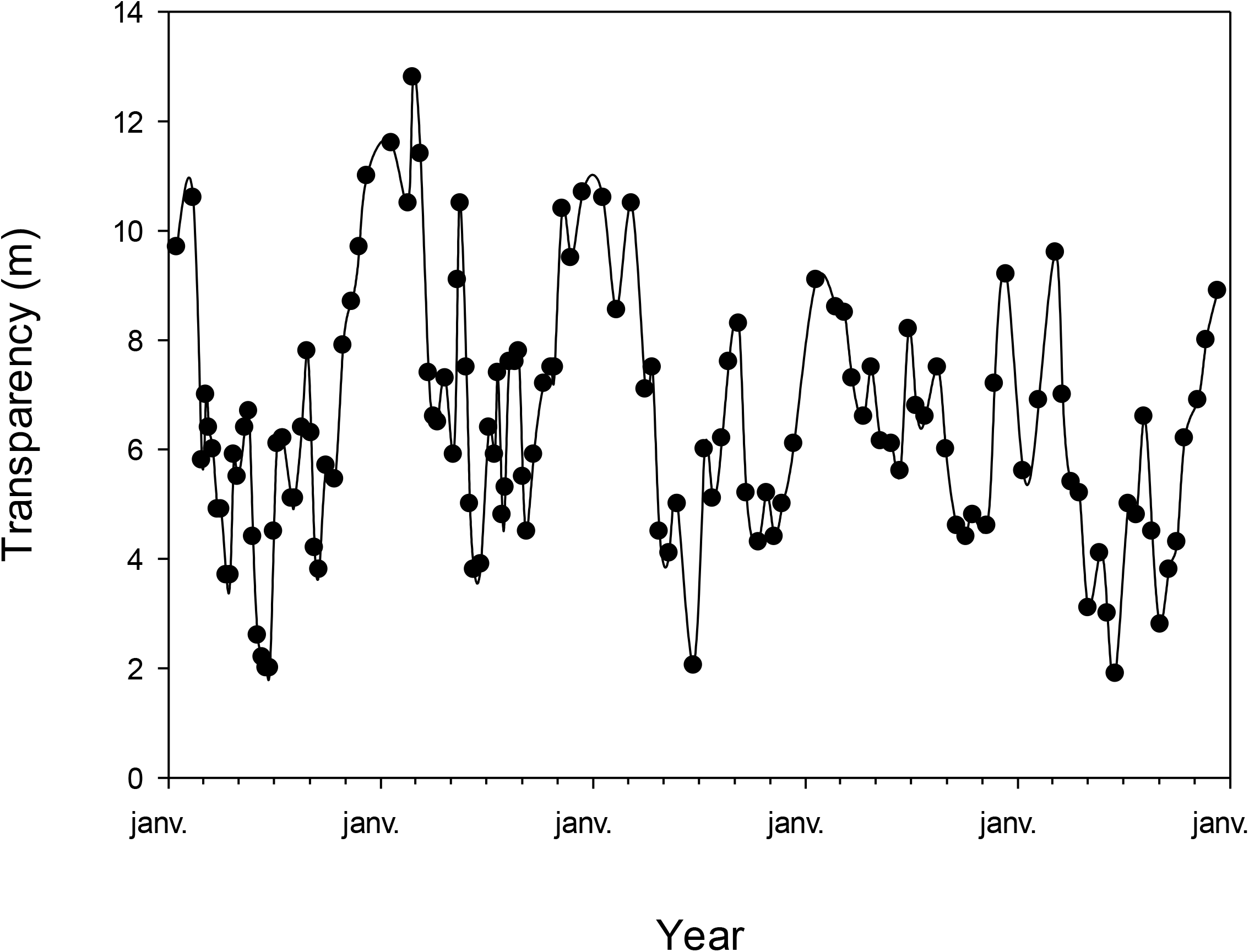
Transparency evolution January 2014 to December 2018.

### Phosphorus input from rivers

From 2010 to 2019, the phosphorus input from the two main lake tributaries (i.e. the Leysse and Sierroz) were distributed differently (not shown). While such input was rather low from 2010 to 2012, it increased significantly at the end of 2012 until the end of 2016. Nutrient loads were also measured to be particularly high when important river floods occurred, such as for example in 2016, June 16^th^ (Fig. 4). During the year 2017, low inflows from the tributaries were observed, whereas the beginning of 2018 was characterised again by high inputs (> 8.6 tons of total-phosphorus and > 0.2 tons of P-PO_4_). In 2019, nutrient loads from the tributaries were still relatively important, especially for PO_4_.

**Fig. 4.**
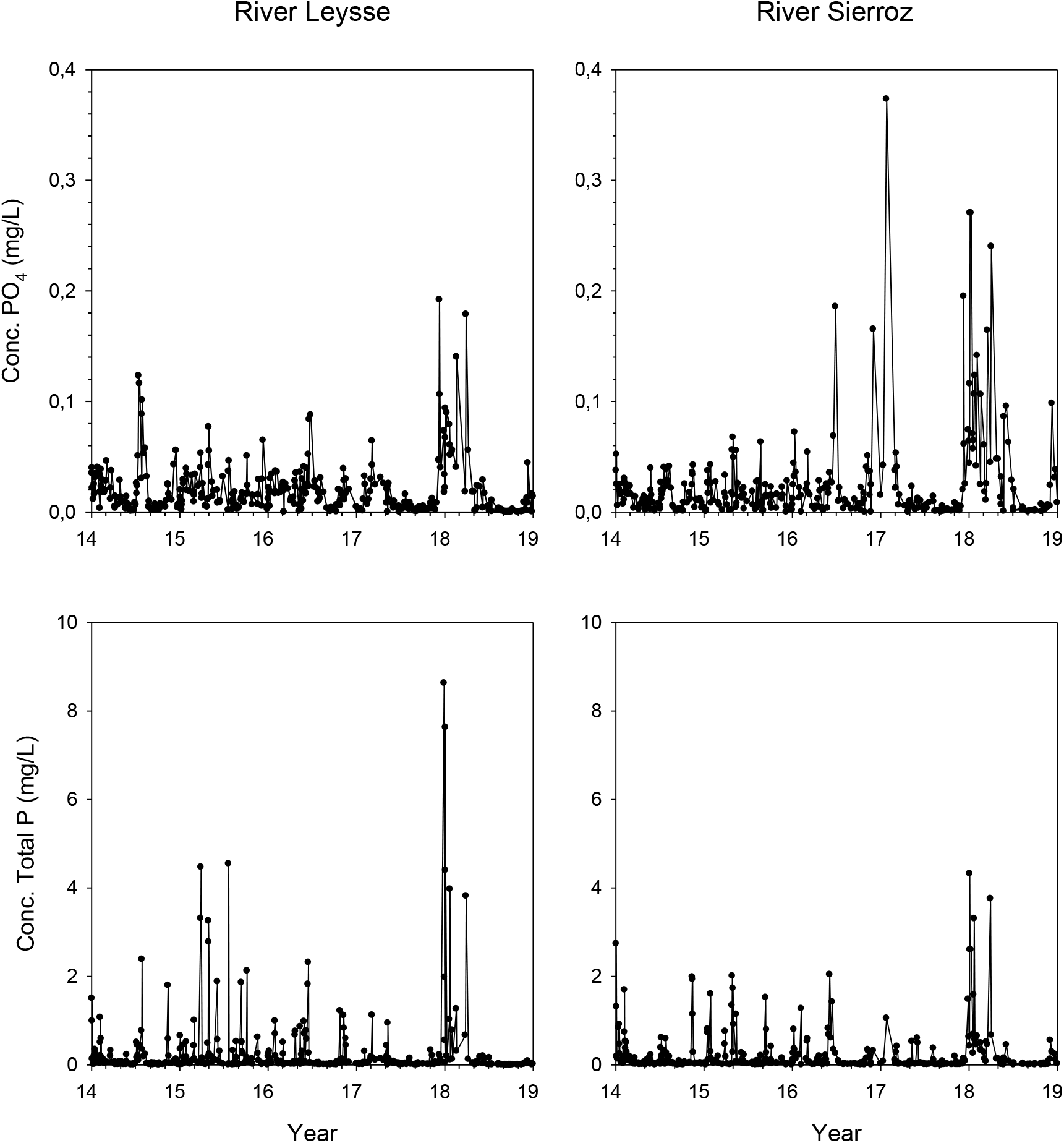
Evolution of total phosphorus and PO_4_ input to the lake from the rivers Leysse and Sierroz from January 2014 to December 2018.

### Phosphorus concentration

From 2010 to 2019, both total phosphorus and orthophosphates concentrations (i.e. the mean values along the water column or in surface waters) decreased regularly. It was observed, however, an increase for the resource for the two winter periods of 2015 and 2016 (Fig. 5 & 6).

**Fig. 5.**
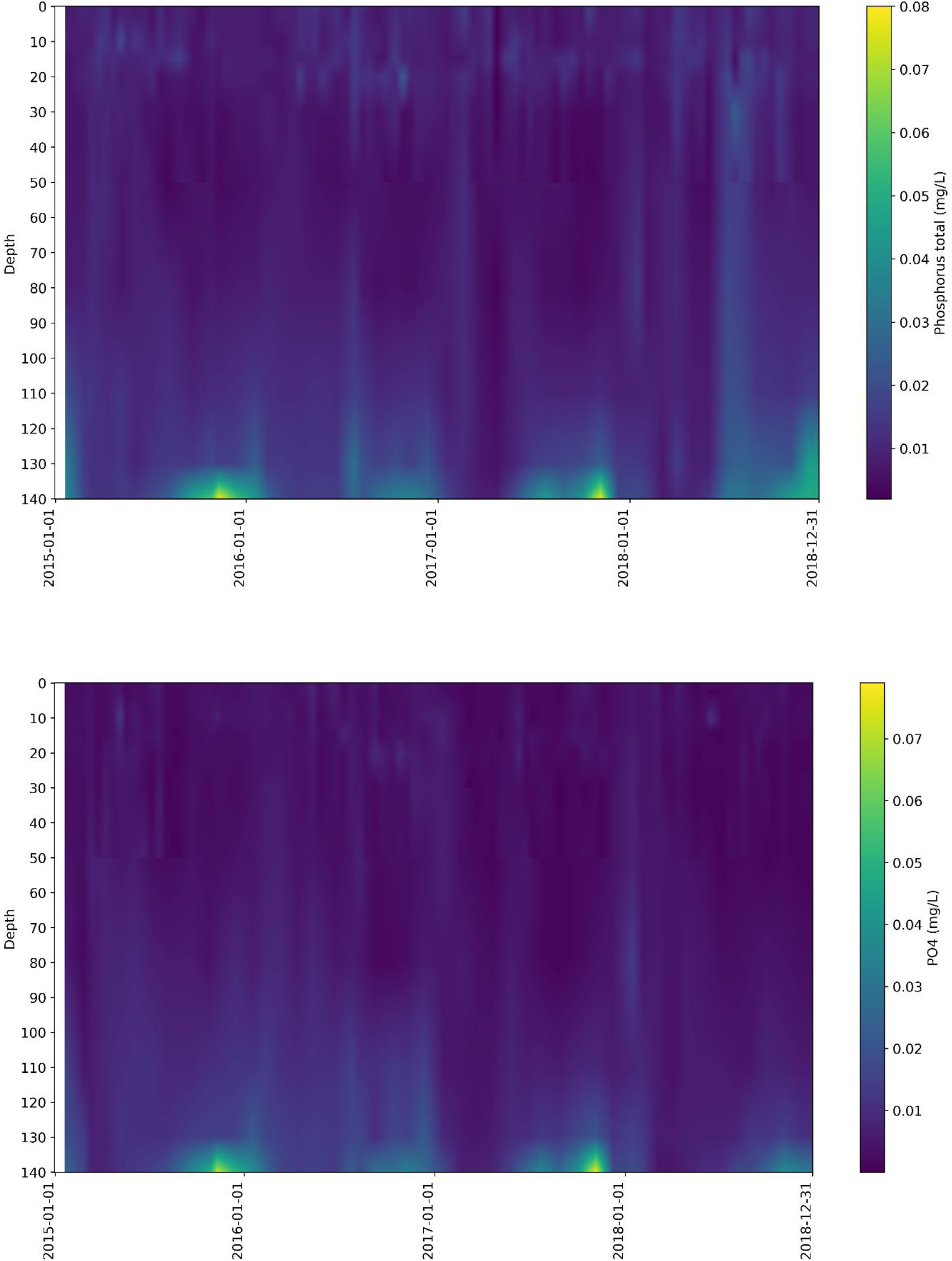
Evolution of total phosphorus and orthophosphate concentrations over the water column, from January 2015 to December 2018

**Fig. 6.**
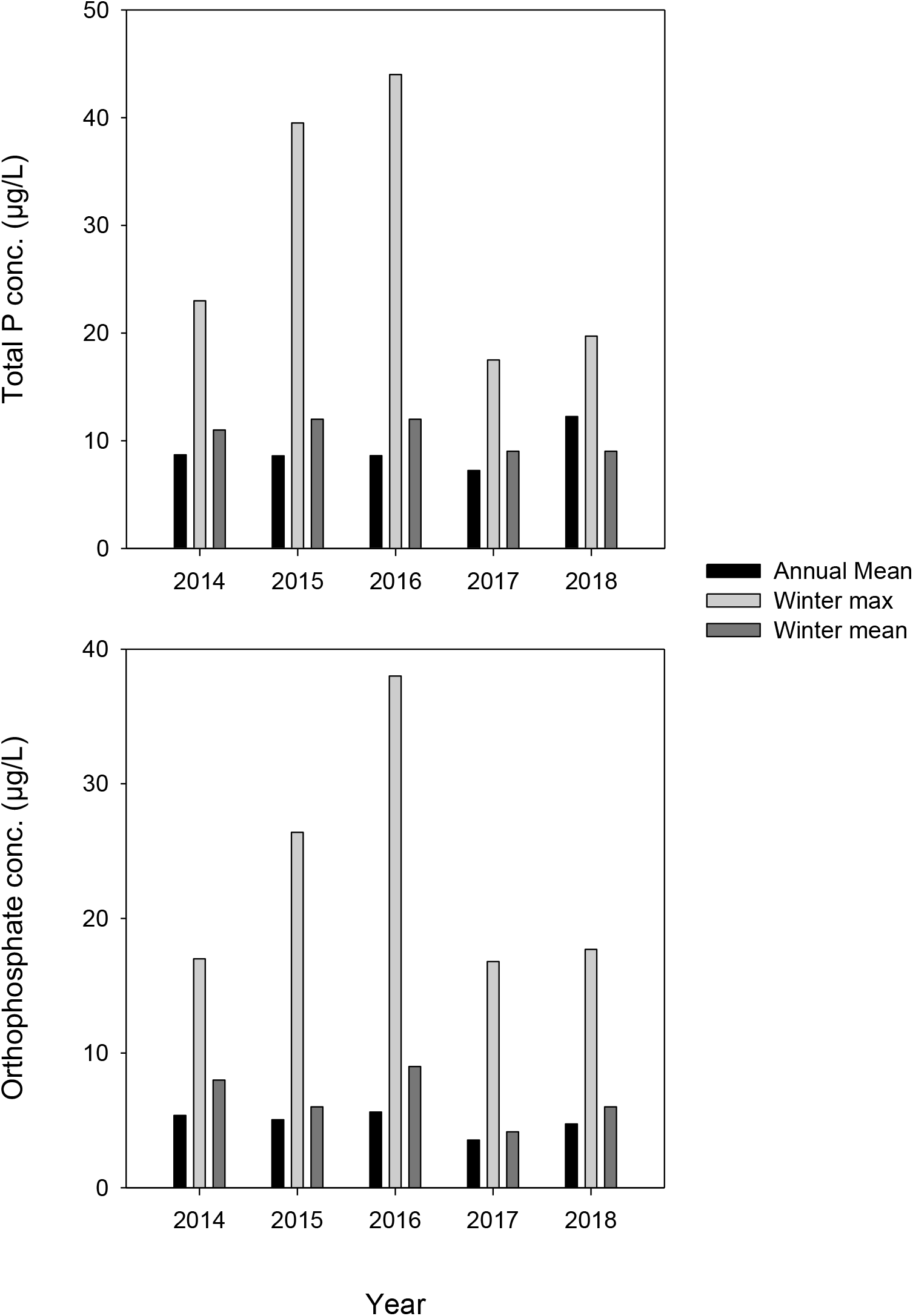
Evolution of winter (max and mean values) and annual (mean value) of total phosphorus and PO_4_ concentrations over the water from 2014 to 2018.

### Water column mixing and stratification

During winter 2016, the water column was poorly homogenised and the partial mixing occurred only from surface to 65 m deep. At mid-April, the maximal dissolved oxygen concentration only reached 6.3 mgO_2_/L. It is noteworthy that the reoxygenation at 140 m was the worst recorded for the last 10 years (not shown). The mixing was also partial in 2017 (despite a colder winter than in 2016) and reached at least 110 m deep. The maximal dissolved oxygen concentration was recorded on February 22^nd^ with 9.2 mgO_2_/L. In 2017, the reoxygenation at 140 m was better than for 2015 and 2016 but stayed significantly lower than for the years 2010 to 2013 (Fig. 7A). At the same time, the stability of the water column, from spring to autumn, was generally important (especially during the summer time). It is noteworthy, however, that mixing and thus the destratification of the water column was relatively important at the end of 2017 (Fig. 7B) while the stability was globally higher for 2016 and 2017 compared to 2015 and 2018 (Fig. 8).

**Fig. 7.**
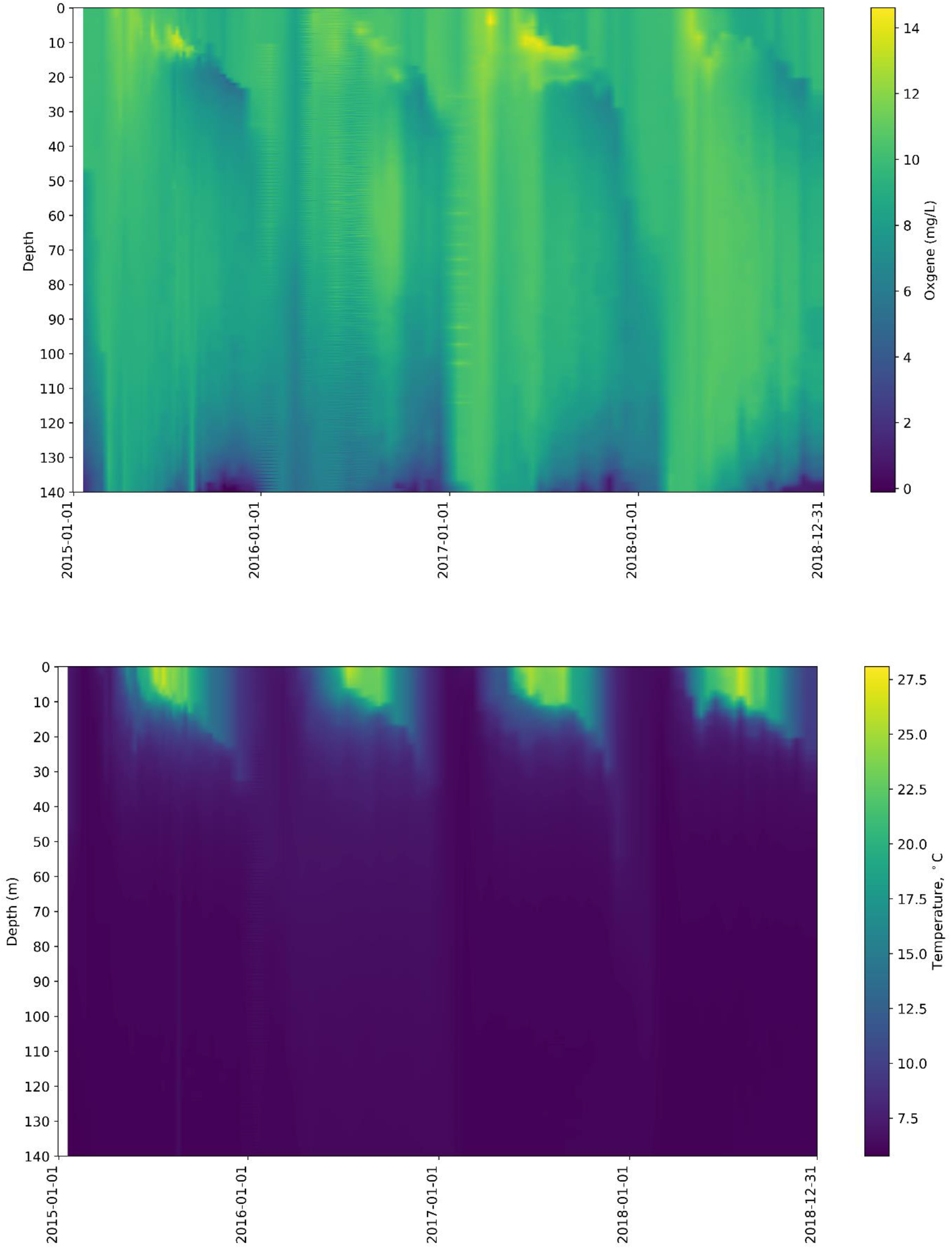
Evolution of dissolved oxygen concentrations and temperature over the water column from 2015 to 2018.

**Fig. 8.**
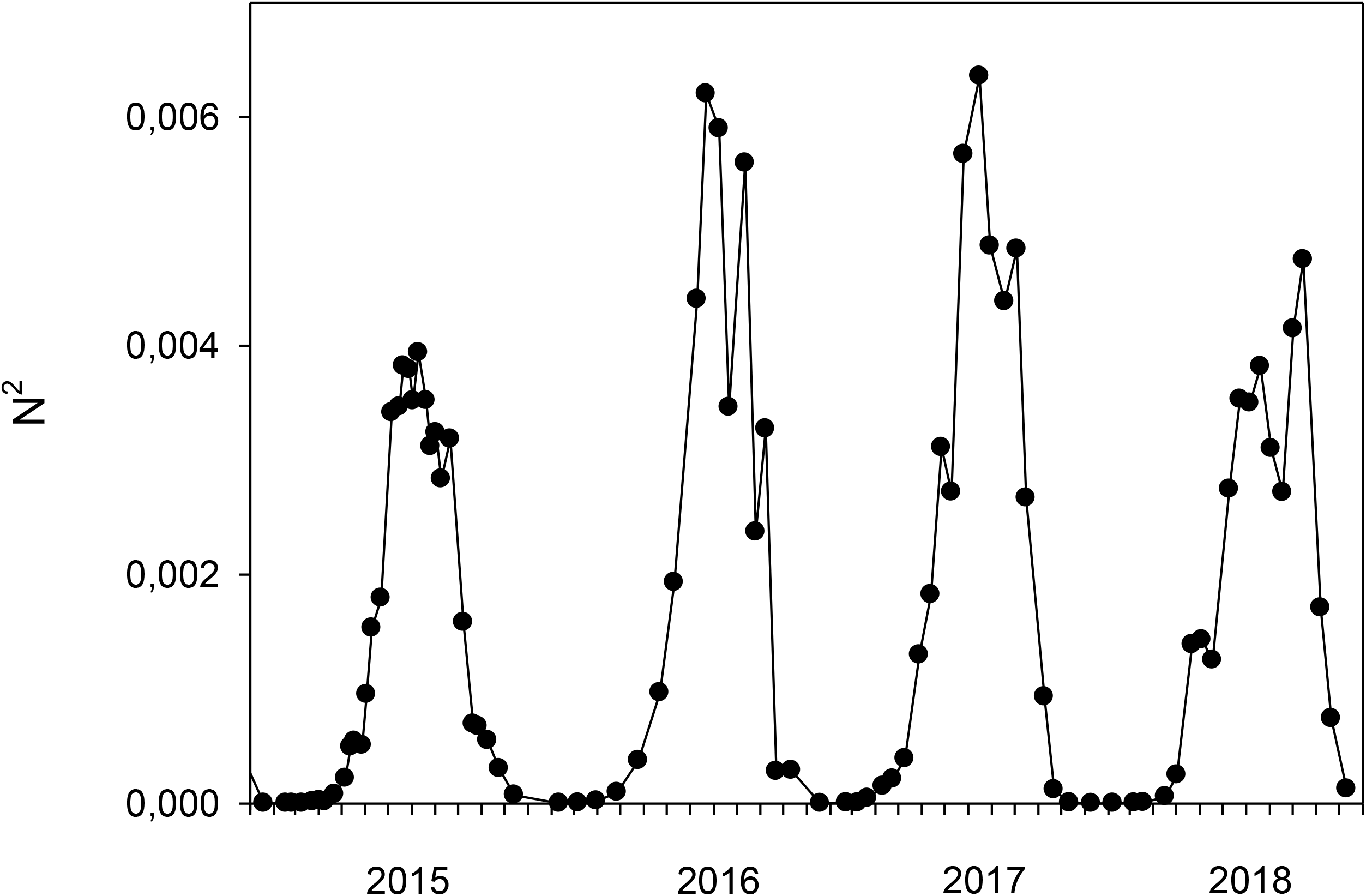
Evolution of the Brunt-Väisälä frequency N^2^ from 2015 to 2018.

### Phyto- and zooplankton

For the last decade, the phytoplankton biomass was higher when *P. rubescens* was recorded during the two years 2016 and 2017, more particularly during the spring season. For the zooplankton, it was usually found a more important density during the summer months (July and August). 2016 and 2017 (with >22,000 ind/mL) were among the years where the zooplanktonic biomass was the highest, especially the proportion of the herbivores (Supplementary Fig. 1).

## Discussion

Amongst aquatic microorganisms, cyanobacteria occupy an important place on Earth because of their historical and ongoing global importance in ecosystem functioning. Moreover, many cyanobacteria are threats since that can generate toxins, especially in inland water bodies. Today, it remains very important to address key issues and highlight key items and new insights regarding these unique organisms. Typically, we still need pieces of information dealing with diversity and functional roles of cyanobacteria, harmful blooms (i.e. determinism, toxin risk, predictive models, management), molecular pathways (including toxin production), abiotic and biotic interactions with cyanobacteria, role of toxin production, as well as about the variety of applications (food supply, socio-economic models). Our study deals with the issue of harmful blooms, more particularly about the attempt to propose some scenarios (even imperfect) about the occurrence of harmful cyanobacterial blooms in lakes (e.g. Anneville et al. 2015, Gallina et al. 2017, Derot et al. 2020). This work makes sense when one knows that future climate scenarios project an increase of such proliferations both in terms of frequency and duration (Paerl & Huisman 2009).

Blooms of *P. rubescens* have been important in Lake Bourget at different periods of the last 3 decades (Jacquet et al. 2005, 2014) and the 2016-2017 episode reminded us that such event may still occur and could impact ecosystem functioning and services (e.g. drinkable water). This study brings new insights on a variety of factors, processes and mechanisms likely to intervene and regulate the blooms of *P. rubescens* in Lake Bourget, which can be viewed and used as a successful model case of ecosystem restoration and reoligotrophication but possibly threatened by cyanobacteria. Indeed, our study highlights that toxic filamentous cyanobacteria such as *P. rubescens* can still proliferate in almost oligotrophic conditions, and thus not only, as a majority of other cyanobacteria, in meso- to eutrophic conditions. We believe that this information is very important for both the scientific community and environmental mangers.

While interesting to the decline of *P. rubescens* in Lake Bourget after a period of important blooms (from 1996 to 2009), Jacquet et al. (2014) reported for the 2009/2010 autumn/winter period that low levels of light, very low phosphorus and P-PO_4_ inputs from tributaries, phosphorus absence or very low concentrations detected in surface waters at the end of the autumn coupled with an absence of partial mixing and consequently possible release of phosphorus from the sediments, altogether these factors and processes largely contributed to the decline of the cyanobacterium. It was suggested, at that time, that an important factor could be the existence during the autumn and/or winter period of a minimal biomass for the cyanobacterium to be able to develop during the following seasons. We tested this last information and found indeed that this autumn/winter inoculum was very important. By simply making average calculations with concentration values at each season, we found indeed that it was necessary for *P. rubescens*, to develop and bloom during the year, to reach a precedent autumn (OND) or winter (either DJF or JFM) threshold above 180 cells/mL (Table 1). Significant positive correlations were clearly found between each season suggesting the importance of the previous months or seasons to explain subsequent blooms later on during the year of *P. rubescens* (Table 2). Thus, by sequencing the years into seasonal periods, and examining the variability in the ecological response of *P. rubescens* to environmental forcing, we show that the successions of events have considerable importance for its development or decline, provided that a minimal initial cell concentration has a determining role for *P. rubescens* dynamics in the following growth season.

**Tab. 1.** Average values calculated for each season of the cell abundance (cells/mL) of *P. rubescens*.

|  | Winter (DJF) | Winter (JFM) | Spring (AMJ) | Summer (JAS) | Autumn (OND) |
| --- | --- | --- | --- | --- | --- |
| 2001 | 7910 | 6196 | 8716 | 9640 | 18521 |
| 2002 | 12600 | 6597 | 719 | 552 | 5028 |
| 2003 | 5429 | 4162 | 1444 | 742 | 45 |
| 2004 | 113 | 169 | 335 | 3662 | 7182 |
| 2005 | 5058 | 1473 | 646 | 5259 | 4328 |
| 2006 | 5796 | 4569 | 1433 | 6216 | 6624 |
| 2007 | 13604 | 11520 | 8187 | 7327 | 5710 |
| 2008 | 5040 | 4107 | 8846 | 18193 | 15958 |
| 2009 | 7504 | 3212 | 1111 | 3659 | 478 |
| 2010 | 32 | 0 | 0 | 4 | 15 |
| 2011 | 22 | 14 | 7 | 2 | 4 |
| 2012 | 12 | 21 | 0 | 0 | 19 |
| 2013 | 0 | 0 | 0 | 0 | 75 |
| 2014 | 71 | 386 | 72 | 0 | 10 |
| 2015 | 0 | 0 | 21 | 75 | 215 |
| 2016 | 193 | 130 | 442 | 4465 | 5775 |
| 2017 | 3827 | 2181 | 1491 | 2739 | 1541 |
| 2018 | 129 | 10 | 17 | 0 | 0 |
| 2019 | 66 | 66 | 66 | 37 | 25 |

**Tab. 2.**
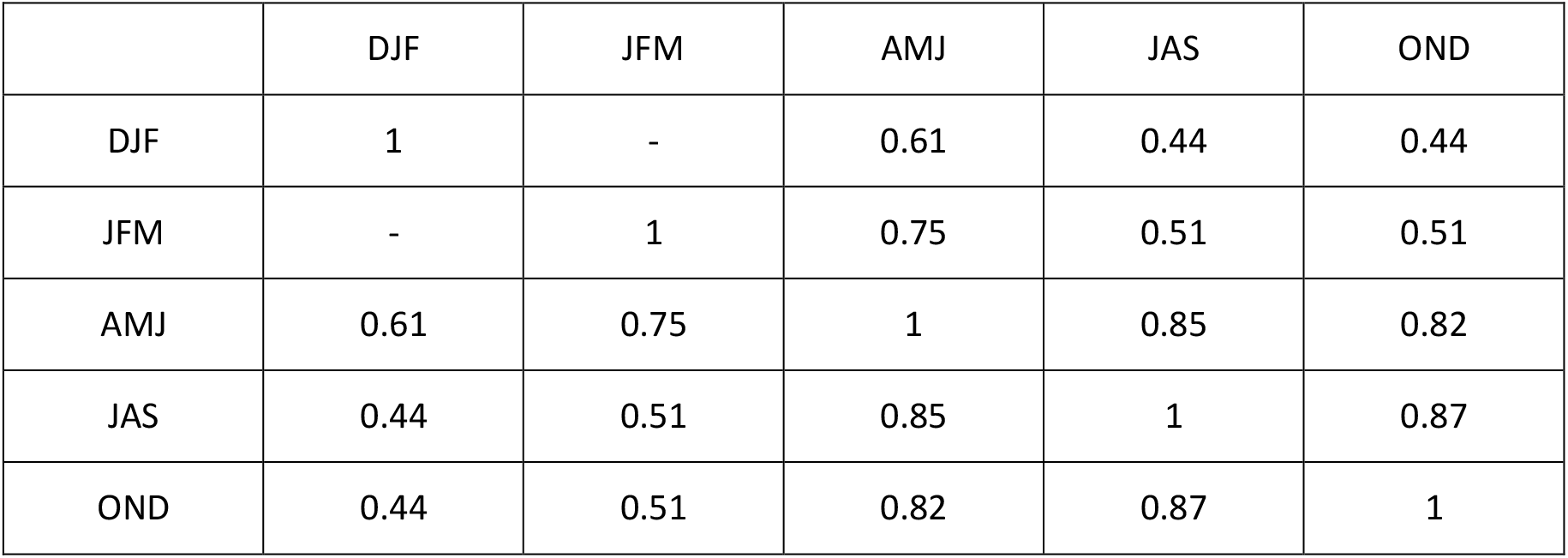
Relationships (r value) between each season for cell abundance (cells/mL) of *P. rubescens* from 2001 to 2020 (n=20).

|  | DJF | JFM | AMJ | JAS | OND |
| --- | --- | --- | --- | --- | --- |
| DJF | 1 | - | 0.61 | 0.44 | 0.44 |
| JFM | - | 1 | 0.75 | 0.51 | 0.51 |
| AMJ | 0.61 | 0.75 | 1 | 0.85 | 0.82 |
| JAS | 0.44 | 0.51 | 0.85 | 1 | 0.87 |
| OND | 0.44 | 0.51 | 0.82 | 0.87 | 1 |

After its disappearance during the 2009/2010 winter period, *P. rubescens* reappeared in the autumn of 2015. During spring 2014, partial water turnover was observed while complete winter mixing occurred between 2010 and 2014. Despite an important sunshine level during this year (2014), these conditions were not sufficient to allow the apparition of the cyanobacterium. It was only at the end of 2015 (November and December), that an inoculum was recorded, linked to favourable conditions such as significant summer inputs of nutrients from tributaries (mainly phosphorus) and a relatively high time of subsequent sunshine and mild winter. Such “warm” conditions and the absence of complete overturn may have reduced the dilution of the cyanobacterium and its growth inhibition or mortality at depth (because of light absence and gas vesicle [that intervene in its buoyancy] collapse under high pressure), not to mention the direct effect of the temperature likely to enhance cell metabolism. Here, we can assume that vertical and lateral transport mechanisms, induced by internal waves or upwelling events, occurred, as already demonstrated by past in Lake Bourget (Cuypers et al. 2011).

Strong inputs of total-phosphorus and P-PO_4_ in June 2016, coupled with significant sunshine during summer and a high stability of the water column, were likely the main factors leading to the important development of *P. rubescens* at depth where it is known to be very competitive. The population ended up being the dominant species starting from summer by forming a dense layer at the metaliminion (due to its low light tolerance and stability requirement) and prevented the growth of other phytoplankters through the reduction of nutrient availability in the upper lit layers. It was in July that the bloom phase was really observed at 15 m depth. With an above-average presence of phosphorus at this period, a high transparency, as well as a well-established stability of the water column and possibly low predation pressure until September, conditions were clearly favourable to allow this new development of *P. rubescens*.

The autumn of 2016 was characterised by a decrease in sunshine and the arrival of autumn gust of wind. Despite a general and progressive decrease in cell density, the search for light by migration was probably the reason of the rise of *P. rubescens* cells to the surface. But the destabilization of the water column due to the autumn wind may have a negative impact on the presence of the cyanobacterium on the surface. Indeed, numerous wind gusts exceeding 30 to 40 km/h, some reaching more than 100 km/h, were recorded during this period.

The decrease in sunshine accompanied by a reduction in transparency, and low nutrient inputs from tributaries during the autumn/winter period of 2016/2017, contributed to the decrease of the cyanobacterium, but without total disappearance. We observed that P could come from the bottom of the lake, because the mild winter of 2016/2017 did not allow a complete water column turnover, and hypoxic conditions at depth were clearly observed, a phenomenon likely to allow the release of phosphorus into the water column. Additionally, the sunshine stayed sufficiently high to allow *P. rubescens* to maintain and grow during the winter, since concentrations were about 2,600 cell/mL on average.

Then, from spring 2017, low phosphorus and P-PO_4_ inputs from tributaries were recorded, but the partial reversal of the water column confirmed at the end of March, and phosphorus gradually moved up from deep sediments, what have compensated this lack. These favourable nutrient conditions, the return of sunshine and a transparency that was maintained at good levels during the spring/summer, allowed the winter inoculum to maintain and be the cause of a summer bloom at depth (above 20 m in spring and early summer) with a well-marked presence between 15 and 30 m. P. *rubescens* beginning from a relatively high concertation, the population could end up being the dominant species starting from summer by forming a dense layer in the metalimnion (due to its low light tolerance and stability requirement) and prevent the growth of other phytoplankters through the reduction of nutrient availability in the upper lit layers. Such competitive exclusion facilitated by the priority effect in which the species with higher initial concentrations outcompete the competitors (characterised by lower initial concentrations) by making the abiotic environment inhabitable has been shown elsewhere and for other species (e.g. Tapolczai et al. 2014).

From September 2017, *P. rubescens* was present in surface, and it was correlated with a decrease in transparency and an important decrease in the sunshine. A reduction in resources with an important water column destratification (favouring dilution of the population) probably contributed to the “disappearance” of *P. rubescens* from January 2018. Despite “record” inputs from the tributaries from January to May, no reappearance occurred, likely due the absence of the inoculum. The decrease in transparency and low spring sunshine probably also contributed to the end of the cycle initiated in winter 2015. While we did not measure it directly here, predation by zooplankton could have also been important as shown by Jacquet et al. (2014) since metazoan feeders can, in some situations (i.e. low growth, size reduction, toxin absence or weak concentrations), impact *P. rubescens* significantly (Oberhaus et al. 2007, Perga et al. 2013, Jacquet et al. 2014, Scharzenberger et al. 2020).

After 2018, the lack of an inoculum, and a sunshine average particularly low prevented *P. rubescens* to develop again. The absence of the cyanobacterium was also confirmed in 2019 and until now (end of 2020). This agrees with previous results (Jacquet et al. 2014), highlighting again that the decline of filamentous and/or colonial cyanobacteria blooms are firstly attributed to phosphate limitation (Walve and Larson 2007) despite the capacity of *Planktothrix* to excrete alkaline phosphatases, allowing for the use of dissolved organic phosphorus when phosphate is depleted (Feuillade et al. 1990).

At last, it is noteworthy that filamentous cyanobacteria can also produce bio-active compounds with antibacterial properties for instance so that such chemical interactions in microbial communities could also play an important role in facilitating the development of the cyanobacterium while preventing the others (Mazur-Marzec et al. 2013, Legrand et al. 2003). We conducted experiments and found indeed that allelopathic effects could be induced by extracts from *P. rubescens* on other phytoplankters (Oberhaus et al. 2008, Chiapusio et al. unpublished).

## Conclusion

Beyond its role as an indicator of environmental degradation and/or major ecological changes occurring in Lake Bourget, bloom forming *P. rubescens* may constitute a serious threat for the lake’s functioning (by inhibiting a part of the matter and energy transfer through the food webs, because of chemical alterations of the water) and ecosystem services (e.g. animal kills, health hazards for humans via drinking water, consumption of fish, recreational use). *P. rubescens* is very opportunistic and can still develop and bloom following environmental shifts inducing favourable conditions for its growth and development. Our results suggest that (i) when meso- to moderately eutrophic conditions are encountered, the proliferation of *P. rubescens* remains possible (Jacquet et al. 2005, Dokulill & Teubner 2012) and (ii) confirm that the success of *P. rubescens* can be attributed to a combination of physico-chemical factors and ecological processes (e.g. Jacquet et al. 2005, 2014, Posch et al. 2012). This study merely suggests that future environmental conditions (extreme events and runoff, temperature increase, deoxygenation and P release from the bottom) may potentially provide conditions for the development and bloom of the cyanobacterium in Lake Bourget, while the latter is oligotrophic. Associated to the counting of a threshold of filaments during the autumn/winter period the year before the bloom (*e*.*g*. >180 cells/mL) and relatively warm winter conditions known to be an important trigger to serve at maintaining the population at a level where it will be possible to proliferate (e.g. Jacquet et al. 2014, Anneville et al 2015, Gallina et al. 2017, Kerimoglu et al. 2017), a simple alert can be imagined and is proposed in Figure 9. Important questions remain: Where did the inoculum come from? Are lateral transport mechanisms induced for instance by internal waves and upwelling events, which occur frequently in Lake Bourget (Cuypers et al. 2011), important drivers for population development? Were filaments present but not detected because of methodical bias due to sampling, sample preparation and/or counting? Or were these filaments located somewhere else in the lake? Etc… In the future, to be able to predict a potential new development of *P. rubescens*, it could be useful to propose a 3D model of the dynamics and distribution of this cyanobacterium, in relation to different factors, themselves linked to the growth of this such particular species.

**Fig. 9.**
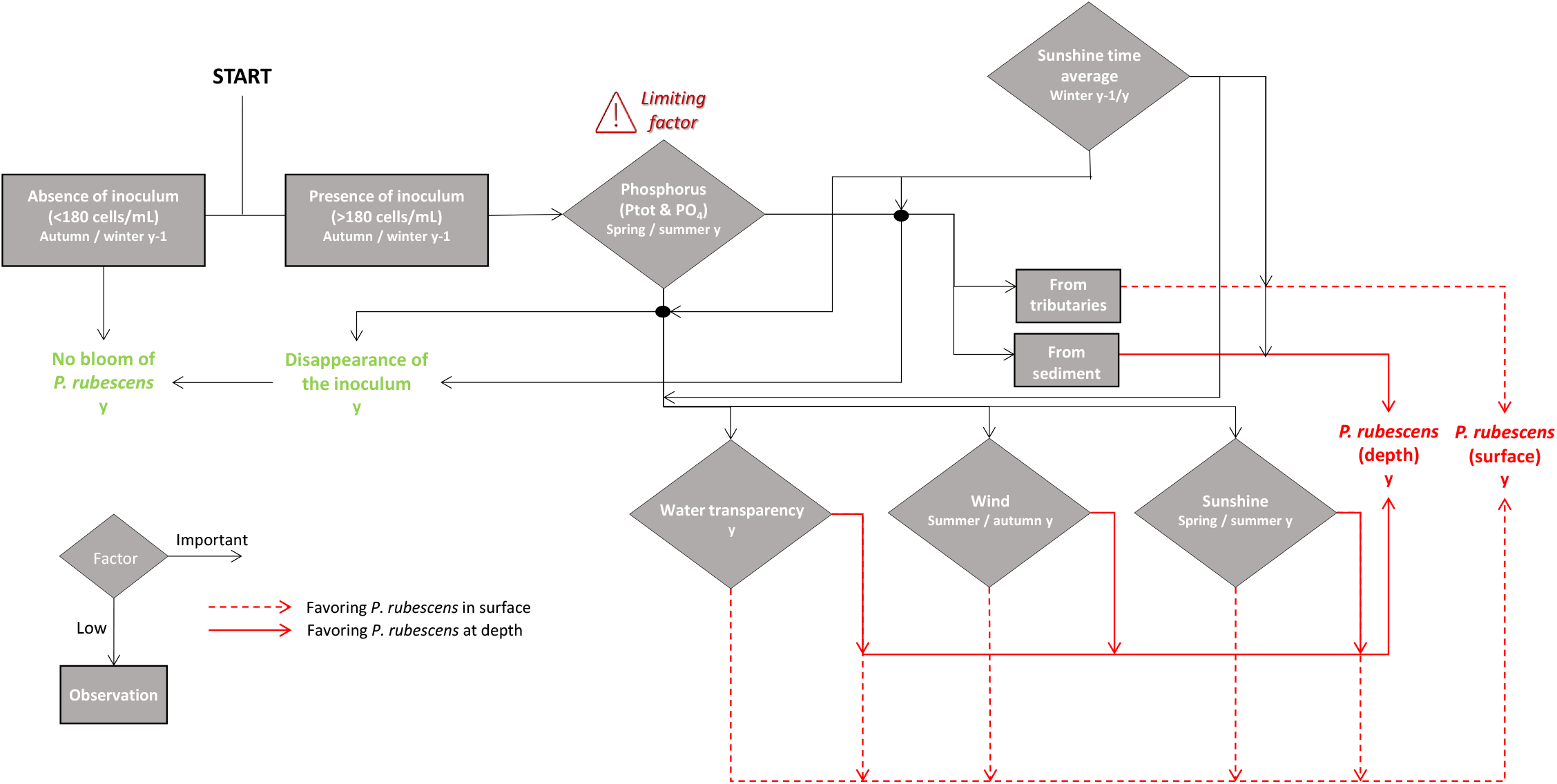
*P. rubescens* is likely to bloom in Lake Bourget following a conjunction of factors and processes. Firstly, the presence of an autumn/winter innoculum of the cyanobacterium the year before is important. Secondly, the phosphorus resource remains a key issue and severe limitation in winter/spring of the year can prevent the bloom whereas presence or input (i.e. P already present in surface waters of the lake or coming from tributaries or from the sediment) during the winter/spring of the year can be crucial. Moreover, a relatively high level of winter irradiance and/or water transparency may clearly favor the initial development of *P. rubescens* merely at depth (i.e. between 15 and 25 m).

## Acknowledgements

We are grateful to the technicians who make possible to obtain a large amount of data of high quality (© OLA-IS, AnaEE-France, INRAE of Thonon-les-Bains, CISALB [Rimet et al. https://doi.org/10.4081/jlimnol.2020.1944]) and Orlane Anneville for her critical reading of a former version of this article.

---

**Supplementary Fig. 1.**
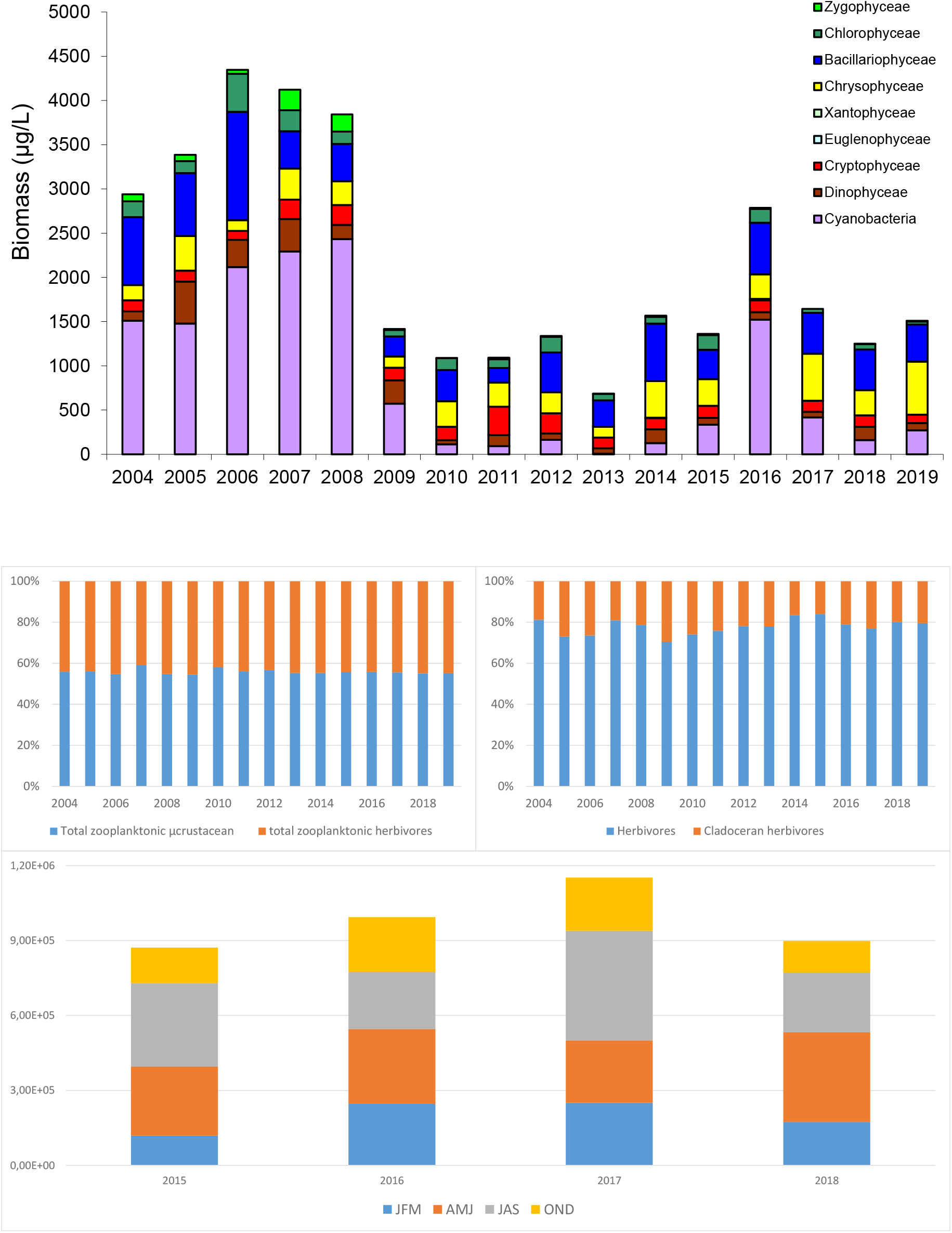
Evolution of the phytoplanktonic biomass (A) using the main classes and the (B) zooplankton using the proportion of the total microcrustacean zooplankton, herbivores and herbivore cladocerans and between 2004 and 2019. A zoom is also proposed for years 2015 to 2018 for the herbivores at the different season.

## Literature cited

Afnor EN 15204. 2006. Water quality - Guidance standard on the enumeration of phytoplankton using inverted microscopy (Utermöhl technique). Afnor: 1–39.

Anagnostidis, K., and J. Komarek. 1988. Modern approach to the classification system of cyanophytes. 3 – Oscillatoriales. Archives of Hydrobiology 80:327–472.

Bright, D. I., and A. E. Walsby. 2000. The daily integral of growth by Planktothrix rubescens calculated from growth rate in culture and irradiance in Lake Zurich. New phytologist 146:301–316.

Briand, J.-F., S. Jacquet, C. Flinois, C. Avois-Jacquet, C. Maisonnette, B. Leberre, and J.-F. Humbert. 2005. Variations in the microcystins production of Planktothrix rubescens (cyanobacteria) assessed by a four-years in situ survey of Lac du Bourget (France) and by laboratory experiments. Microbial Ecology 50:418–428.

Capo, E., D. Debroas, F. Arnaud, T. Guillemot, V. Bichet, L. Millet, E. Gauthier, and I. Domaizon. 2016. Long-term dynamics in microbial eukaryotes communities: A palaeolimnological view based on sedimentary DNAMolecular Ecology 25 (23):5925–5943.

Capo, E., D. Debroas, F. Arnaud, M.E. Perga, C. Chardon, and I. Domaizon. 2017. Tracking a century of changes in microbial eukaryotic diversity in lakes driven by nutrient enrichment and climate warming. Environmental microbiology 19 (7):2873–2892.

Derot, J., H. Yajima, and S . Jacquet. 2020. Advances in forecasting harmful algal blooms using machine learning models: A case study with Planktothrix rubescens in Lake Geneva. Harmful Algae 99:101906.

Dokulil, M.T., and K. Teubner. 2012. Deep living Planktothrix rubescens modulated by environmental constraints and climate forcing. Hydrobiologia 698:29–46.

Fastner, J., M. Erhard, W. W. Carmichael, F. Sun, K. L. Rinehart, H. Rönicke and I. Chorus. 1999. Characterization and diversity of microcystins in natural blooms and strains of the genera Microcystis and Planktothrix from German freshwaters. Archives of Hydrobiology 145:147–163.

Feuillade, J. 1994. The cyanobacterium (blue-green algae) Oscillatoria rubescens D.C. Arch. Hydrobiol. Beih. Eregbn. Limnol. 41:77–93.

Frossard, V., N. Grandrémy, F. Arthaud, J. Guillard, and S. Jacquet. Evidence of ecological shifts in fresh water ecosystems: A case study with Lake Bourget (France).

Gallina, N., M. Beniston, and S. Jacquet. 2017. Estimating future cyanobacterial occurrence and importance in lakes: a case study with Planktothrix rubescens in Lake Geneva. Aquat Sci 79:249–263.

Jacquet, S., J.-F. Briand, C. Leboulanger, C. Avois-Jacquet, L. Oberhaus, B. Tassin, B. Vinçon-Leite, G. Paolini, J.-C. Druart, O. Anneville and J.-F. Humbert. 2005. The proliferation of the toxic cyanobacterium Planktothrix rubescens following restoration of the largest natural French lake (Lac du Bourget). Harmful Algae 4:651–672.

Jacquet, S., O. Kerimoglu, F. Rimet, G. Paolini and O. Anneville. 2014. Cyanobacterial bloom termination: the disappearance of Planktothrix rubescens from Lake Bourget after restoration. Freshwater Biology59:2472-2489.

Jacquet, S., F. Arthaud, D. Barbet, C. Barbier, S. Cachera, L. Crépin, L. Espinat, C. Goulon, J. Guillard, V. Hamelet, J.C. Hustache, L. Laine, R. Lambert, A. Miquet, J. Neasat, G. Paolini, P. Perney, P. Quétin, F. Rimet, SF Rivera Rocabado. 2017. Suivi environnemental des eaux du lac du Bourget pour l’année 2016. Rapport INRA-CISALB-CALB, 211 pages.

Jenny, J.P., and 40 co-authors 2020. Scientists’ Warning to Humanity: Rapid degradation of the world’s large lakes. Journal of Great Lakes Research 46:686–702.

Kerimoglu, O., S. Jacquet, B. Vinçon-Leite, B. Lemaire, F. Rimet, F. Soulignac, D. Trévisan, O. Anneville. 2017. Modelling the plankton groups of the deep, peri-alpine Lake Bourget. Ecological Modeling 359:415–433.

Kurmayer, R., J.F. Blom, L. Deng, and J. Pernthaler. 2015. Integrating phylogeny, geographic niche partitioning and secondary metabolite synthesis in bloom-forming Planktothrix. Isme Journal 9:909–921.

Leboulanger, C., U. Dorigo, S. Jacquet, B. LeBerre, G. Paolini, and J.-F. Humbert. 2002. Application of a submersible spectrofluorometer for rapid monitoring of freshwater cyanobacterial blooms: a case study. Aquatic Microbial Ecology 30:83–89.

Mazur-Marzec, H. and 10 co-authors. 2013. Occurrence of cyanobacteria and cyanotoxins in the Southern Baltic Proper. Filamentous cyanobacteria vs. single-celled picocyanobacteria. Hydrobiologia 701(1):235–252.

Micheletti, S., Schanz, F. & Walsby, A. E. 1998. The daily integral of photosynthesis by Planktothrix rubescens during summer stratification and autumnal mixing in Lake Zürich. New Phytologist 138: 233–249.

Oberhaus, L., M. Gelinas, B. Pinel-Alloul, A. Ghadouani, and J.-F. Humbert. 2007. Grazing of tow toxic Planktothrix species by Daphnia pulicaria: potential for bloom control and toxin transfer of microcystins. Journal of Plankton Research 29:827–838.

Oberhaus, L., J.-F. Briand, and J.-F. Humbert. 2008. Allelopathic growth inhibition by the toxic, bloom-forming cyanobacterium Planktothrix rubescens. FEMS Microbiology Ecology 66:243–249.

Posch, T., O. Koster, M.M. Salcher, and J. Pernthaler. 2012. Harmful filamentous cyanobacteria favoured by reduced water turnover with lake warming. Nature Climate Change 2:809–813.

Reynolds, C., V. Huszar, C. Kruk, L. Naselli-Flores, and S. Melo. 2002. Towards a functional classification of the freshwater phytoplankton. Journal of Plankton Research 24:417–428.

Rimet, F., and 20 co-authors. 2020. The Observatory on LAkes (OLA) database: Sixty years of environmental data accessible to the public. Journal of Limnology DOI: 10.4081/jlimnol.2020.1944.

Rimet, F., and J.-C. Druart. 2018. A trait database for phytoplankton of temperate lakes. Annales de Limnologie - International Journal of Limnology 54:18.

Savichtcheva, O., D. Debroas, M.E. Perga, F. Arnaud, C. Villar, E. Lyautey, A. Kirkham, C. Chardon, B. Alric, and I. Domaizon. 2015. Effects of nutrients and warming on Planktothrix dynamics and diversity: a palaeolimnological view based on sedimentary DNA and RNA. Freshwater Biology 60:31–49.

Schwarzenberger, A., R. Kurmayer, and D. Martin-Creuzburg. 2020. Toward disentangling the multiple nutritional constraints imposed by Planktothrix: the significance of harmful secondary metabolites and sterol limitation. Frontiers in Microbiology 11:586120.

Sotton, B., O. Anneville, S. Cadel-Six, I. Domaizon, S. Krys, and J. Guillard. 2011. Spatial match between Planktothrix rubescens and whitefish in a mesotrophic peri-alpine lake: evidence of toxins accumulation. Harmful Algae 10:749–758.

Sotton, B., J. Guillard, S. Bony, A. Devaux, I. Domaizon, N. Givaudan, H. Huet, and O. Anneville. 2012. Impact of toxic cyanobacterial blooms on Eurasian perch (Perca fluviatilis): experimental approaches and in situ observations in a peri-alpine lake. PLoS ONE 7(12):e52243.

Strickland, J. D. H, and T. R. Parsons 1972. A practical handbook of seawater analysis. 2nd Ed. Bull. Fish. Res. Bd. Canada 167: 311 p.

Uthermöhl, H. 1958. Zur Vervollkommung der quantitativen phytoplankton-methodik. Mitt. Int. Ver. Limnol. 9, 38 p.

Vinçon-Leite, B., B. Tassin, and J.-C. Druart. 2002. Phytoplankton variability in Lake Bourget: Phytoplankton dynamics and meteorology. Lakes & Reservoirs: Research and Management 7:93–102.

Walve, J., and U. Larsson. 2007. Blooms of Baltic Sea Aphanizomenon sp. (Cyanobacteria) collapse after internal phosphorus depletion. Aquatic Microbial Ecology 49:57–69.

Zotina, T., O. Koster, and F. Juttner. 2003. Photoheterotrophy and light-dependent uptake of inorganic and organic nitrogenous compounds by Planktothrix rubescens under low irradiance. Freshwater Biology 48:1859–1872.

